# DIALib: an automated ion library generator for data independent acquisition mass spectrometry analysis of peptides and glycopeptides

**DOI:** 10.1101/696518

**Authors:** Toan K. Phung, Lucia F Zacchi, Benjamin L. Schulz

## Abstract

Data Independent Acquisition (DIA) Mass Spectrometry (MS) workflows allow unbiased measurement of all detectable peptides from complex proteomes, but require ion libraries for interrogation of peptides of interest. These DIA ion libraries can be theoretical or built from peptide identification data from Data Dependent Acquisition (DDA) MS workflows. However, DDA libraries derived from empirical data rely on confident peptide identification, which can be challenging for peptides carrying complex post-translational modifications. Here, we present DIALib, software to automate the construction of peptide and glycopeptide Data Independent Acquisition ion Libraries. We show that DIALib theoretical ion libraries can identify and measure diverse *N*- and *O*-glycopeptides from yeast and mammalian glycoproteins without prior knowledge of the glycan structures present. We present proof-of-principle data from a moderately complex yeast cell wall glycoproteome and a simple mixture of mammalian glycoproteins. We also show that DIALib libraries consisting only of glycan oxonium ions can quickly and easily provide a global compositional glycosylation profile of the detectable “oxoniome” of glycoproteomes. DIALib will help enable DIA glycoproteomics as a complementary analytical approach to DDA glycoproteomics.

## Introduction

Key challenges in bottom-up mass spectrometry (MS) experiments are peptide identification and quantification. The wide range of protein abundance in complex samples and the presence of post-translational modifications reduce the efficiency of both peptide identification and quantification. One of the most common post-translational modifications in proteins is glycosylation [1, 2]. Protein glycosylation is also particularly difficult to analyse by MS, due to the natural site-specific heterogeneity in glycan structure and occupancy [3]. Site specific variation in the structure or occupancy of glycans can lead to major changes in glycoprotein stability, folding, and function in physiological, pathological, and biotechnological situations [3–10]. Driven by the importance of glycans in biology, substantial progress has been made in analytical LC-MS/MS methods to identify and quantify glycan heterogeneity in complex samples [11, 12].

Data Independent Acquisition (DIA) is a major recent development in mass spectrometry proteomics. In DIA, the decision to fragment a particular precursor is made not based on the precursor’s intensity, but rather on its presence within predefined *m/z* windows [13, 14]. One implementation of DIA is sequential window acquisition of all theoretical mass spectra (SWATH) [13]. DIA workflows are powerful for unbiased measurement of the abundance of all detectable peptides from complex proteomes. However, DIA workflows depend on the construction of ion libraries that allow measurement of specific precursor and fragment ion *m/z* pairs (a.k.a. transitions) with specific RTs. These DIA ion libraries are routinely built from peptide identification data obtained through Data Dependent Acquisition (DDA) workflows. The main limitation of these empirical DDA-derived libraries is that the peptides identified passed the intensity cut-off requirements for DDA fragmentation and could be identified using database searching software. These two requirements limit the potential of DIA proteomics, especially for investigation of post-translationally modified peptides. One strategy to circumvent this limitation is to build theoretical ion libraries that include all the desired transitions that would result from fragmentation of peptides modified with post-translational modifications of interest.

We previously used a SWATH DIA approach to measure relative differences in glycan occupancy and structure in yeast cell wall proteins as a consequence of defects in the *N*-glycosylation machinery [5]. This approach used a manually constructed ion library composed of *b* and *y* peptide ions and glycan oxonium ions. We observed that while all mutants showed distinct and significant differences in glycan occupancy compared to the wild-type strain, the mannosyltransferase mutants (*alg3Δ*, *alg9Δ*, and *alg12Δ*) also displayed significant changes in glycan structure due to defects in the addition of mannose residues to the glycan branches [5]. However, because the manually curated library depended on our ability to manually identify glycopeptides with multiple glycoforms in the DDA data, the ion library generated most likely underrepresented the diversity and heterogeneity of the yeast cell wall glycoproteome.

An alternative way to obtain a more comprehensive representation of glycoproteome diversity is to measure glycopeptides using Y ions present in MS/MS fragmentation spectra. Y ions are high intensity ions produced during glycopeptide fragmentation with CID/HCD, and consist of the entire precursor peptide sequence with 0, 1, or more monosaccharide residues attached to the glycosylation site (termed Y0, Y1, Y2, etc) [15]. Y ions are a common fragment ion to all glycoforms of the same glycopeptide, independent of glycan structure, and are thus an ideal fragment ion to use to detect and measure glycan structural heterogeneity without prior information on glycan structures or glycopeptide precursor *m/z* [15]. This approach has been shown to be effective in identifying common and rare glycoforms in a glycopeptide-enriched serum sample [15].

The manual construction of spectral libraries can be tedious and time consuming. Here, we present DIALib, software to automate the construction of peptide and glycopeptide **D**ata **I**ndependent **A**cquisition ion **Lib**raries, for use in DIA analyses with Peakview (SCIEX). We show that DIALib theoretical Y libraries can identify and measure *N*- and *O*-glycopeptides with and without prior knowledge of the glycoforms present and of their retention times, and we discuss the utility of complementing Y libraries with *b* and *y* peptide fragment ions. We also show that DIALib libraries consisting only of glycan oxonium ions can quickly and easily provide a global glycosylation profile of DIA samples.

## Methods

### Library construction

DIALib is software that can generate customized ion libraries to use in Peakview (SCIEX). https://github.com/bschulzlab/DIALib. Briefly, the software processes the input protein or peptide sequences to generate ion libraries containing peptide *b* and *y* ions, glycopeptide Y ions, and/or oxonium ions of choice, and allows the user to select the retention times (RT) to be used for the peptide(s) and precursor masses. Key features of the software are described in detail below.

#### Input data

The input data for the software is one or more protein or peptide sequences in FASTA format, or Uniprot identifiers (www.uniprot.org). The fasta file or text input is first processed to retrieve protein identification and amino acid sequence. DIALib can perform *in silico* protease digestion of the input sequences, allowing selection of specific peptides of interest for downstream processing and inclusion in the library. The user can select from a range of default settings or input a customized setting of choice, including: a) Modifications (static (e.g. propionamide or carbamidomethyl), variable Y-type (e.g. Y0, Y1, and/or Y2 ions for Hex or HexNAc), and variable non-Y-type (e.g. phosphorylation, sulfation, and carboxylation, among others); b) RT (unique or multiple selection); c) *m/z* windows, corresponding to the Q1 value (normally, the precursor *m/z* values) (unique or multiple selection); d) Maximum ion fragment charge; e) Oxonium ions (unique or multiple selection); and f) Fragment ion types, corresponding to the Q3 values (normally, the fragment ion *m/z* values) (*b*, *y*, or Y). Finally, the user can choose to generate a library that contains multiple RT, or multiple *m/z* windows, or both.

RT selection: the user can select specific RT (if prior RT information is available) or a range of RTs (if no prior RT information is available).

Q1 selection: DIALib allows the user to input specific Q1 values or to select a range of *m/z* windows in which specific transitions will be measured. Multiple Q1s are especially important when no prior information of Q1 is available, or when multiple Q1 for the same peptide are expected due to the presence of PTMs with unpredictable *m/z* (e.g. glycosylation). If multiple Q1s are selected, the value of Q1 that appears in the library is the middle value of the chosen *m/z* range (e.g. if the *m/z* window chosen is 400-425 *m/z*, the ion library will show a Q1 value of 412.5).

Q3 selection: Libraries made of *b* and *y* ions (*by* libraries) do not contain the *b1*, *y1*, and full length *b* ion. It is possible to suggest a stop position within the peptide to limit the *b* or *y* series, and in this way *b* and *y* transitions are only generated until the stop position. It is also possible to select specific *b* and *y* transitions for each peptide. The type and number of transitions used in the library can be modified for each peptide. In the case in which the library contains *b*, *y*, and either Y ions or variable post-translational modifications, there is an option to generate *b* and *y* transitions that correspond to either the modified or unmodified query peptide (i.e. they contain or not the mass of the post-translational modification that is added to the Y ion series).

#### Library generation

Based on input data and user defined settings, DIALib generates a combination of query peptides with modified sites and single or multiple RTs. The ion library is generated as a text file and contains all the parameters that Peakview requires, including parameters that are fixed for all transitions: relative intensity (set to 1), score (set to 0), prec_y (set to 0), confidence (set to 0.99), and shared (set to FALSE) (Supplementary Table S1).

### Preparation of human serum-derived Immunoglobulin G

Purified human IgG (I4506, Sigma) was prepared essentially as previously described [16, 17]. 5 µg of IgGs were denatured and reduced in a buffer containing 6 M guanidine hydrochloride, 50 mM Tris pH 8, and 10 mM dithiothreitol (DTT) for 30 minutes at 30 °C while shaking at 1500 rpm in a MS100 Thermoshaker incubator (LabGene Scientific Ltd.). Reduced cysteines were alkylated by addition of acrylamide to a final concentration of 25 mM followed by 1 h incubation at 30 °C in a thermoshaker at 1500 rpm, and excess acrylamide was then quenched by addition of DTT to a final additional concentration of 5 mM. Proteins were precipitated by addition of 4 volumes of 1:1 methanol/acetone and incubation for 16 h at −20 °C. After centrifugation at 18,000 rcf at room temperature for 10 min, the supernatant was discarded and the precipitated proteins were resuspended in a 50 mM ammonium bicarbonate buffer. Proteins were digested overnight with trypsin (T6567, Sigma) at 37 °C in a thermoshaker at 1500 rpm. Peptides were desalted by ziptipping with C18 Ziptips (ZTC18S960, Millipore).

### Mass Spectrometry analysis

Desalted peptides were analyzed by liquid chromatography electrospray ionization tandem mass spectrometry (LC-ESI-MS/MS) using a Prominence nanoLC system (Shimadzu) and a TripleTof 5600 mass spectrometer with a Nanospray III interface (SCIEX) essentially as described [5, 18]. Using a flow rate of 30 µl/min, samples were desalted on an Agilent C18 trap (0.3 x 5 mm, 5 µm) for 3 min, followed by separation on a Vydac Everest C18 (300 Å, 5 µm, 150 mm x 150 µm) column at a flow rate of 1 µl/min. A gradient of 10-60% buffer B over 45 min where buffer A = 1 % ACN / 0.1% FA and buffer B = 80% ACN / 0.1% FA was used to separate peptides. Gas and voltage settings were adjusted as required. MS TOF scan across 350-1800 *m/z* was performed for 0.5 sec followed by data dependent acquisition (DDA) of up to 20 peptides with intensity greater than 100 counts, across 100-1800 *m/z* (0.05 sec per spectra) using collision energy (CE) of 40 +/− 15 V. For data independent (DIA) analyses, MS scans across 350-1800 *m/z* were performed (0.5 sec), followed by high sensitivity DIA mode with MS/MS scans across 50-1800 *m/z*, using 34 isolation windows of width 26 *m/z*, overlapping by 1 *m/z*, for 0.1 sec, across 400-1250 *m/z*. CE values for SWATH samples were automatically assigned by Analyst software (SCIEX) based on *m/z* mass windows.

### Data analysis

The ion libraries generated using DIALib were used to measure the abundance of glycopeptides with different glycan structures using Peakview (v2.2, SCIEX). The Peakview processing settings employed were, unless specifically noted, as follows: all peptides were imported, all transitions/peptide were used for quantification, the confidence threshold was 99%, the False Discovery Rate (FDR) was 1% (except for the oxoniome quantitation, in which 100% FDR was used), and the error allowed was 75 ppm. The width of the XIC RT window varied depending on the analysis performed. For identifying and quantifying *N*-glycoforms in the yeast cell wall samples [5], an XIC RT window of 25 min, which covered the entire LC gradient, was used with the Y ion library, and an XIC RT window of 2 min was used with the optimized RT *by* ion library. For identifying and quantifying *N*-glycoforms in human IgG samples and *O*-glycoforms in the yeast cell wall samples [5], an XIC RT window of 1 or 2 min was used, respectively. All data was exported from PeakView allowing 100% FDR. To measure site-specific glycan structure, data was first normalized as follows. The signal intensity measurement for all transitions from a given glycopeptide at a particular Q1 and RT was divided by the summed abundance of all detected signal of that glycopeptide for all Q1s at all RTs. The normalized signal intensities obtained from multiple RT and multiple Q1 data was re-arranged using an in-house developed script called reformat_peptide3.py (Supplementary Information) in which the observed RT was rounded up to the nearest integer and then the data was arranged in a matrix of RT *vs m/z* window for each peptide. The yeast cell wall MS analyses were performed as biological triplicates [5], while the IgG sample was n=1. Heatmaps were prepared in Prism v7.00 (GraphPad Software, La Jolla California USA). Statistical analyses were performed using two-tailed *t* test in Excel (Microsoft).

Byonic (Protein Metrics v2.13.17) was used to identify *N*- and *O*-glycopeptides in the human IgG sample and in the yeast cell wall samples. For both searches, Preview (Protein Metrics v2.13.17) was first used to obtain an estimate of the precursor and fragment *m/z* errors in the MS files, a key parameter for subsequent searches in Byonic. The searches in Byonic for *N*-glycans in human IgG were performed with the following settings: Protein database: *Homo sapiens* proteome UP000005640 downloaded from Uniprot on April 20^th^ 2018 (with 20,303 reviewed proteins) with decoys and common contaminants added; Digestion specificity was set to Fully specific and no misscleavages were allowed (to only identify glycoforms of the fully cleaved tryptic version of the IgGs, mimicking the Y ion library); and Precursor and Fragment mass tolerance were set to 20 and 30 ppm, respectively. Propionamide was set as fixed modification. The only modification included was a modified version of the Byonic database of 57 *N*-glycans at the consensus sequence N-X-S/T containing 8 additional glycoforms, set as “Common 1”. Only 1 common modification was allowed per peptide. To identify *N*- and *O*-glycans in the cell wall sample, we searched one DDA sample of the wild-type yeast. Byonic searches were performed with the following settings: 1) Protein database: *Saccharomyces cerevisiae* proteome UP000002311 downloaded from Uniprot on April 20^th^ 2018 (with 6049 reviewed proteins) with decoys added; Digestion specificity was set to Fully specific and one misscleavage was allowed; Precursor and Fragment mass tolerance were set to 40 ppm and 1 Da, respectively. Propionamide was set as a fixed modification. The modifications allowed included a homemade database of yeast *N*-glycans at the consensus sequence N-X-S/T, set as “Common 1”, or *O*-glycans at S/T set as “Rare 1”. The *N*-glycan database contained structures with 2 HexNAc and 1 to 15 Hex, and the *O*-glycan database contained structures with 1-12 Hexoses. Only 1 common and 1 rare modifications were allowed per peptide.

## Results

### Automated construction of theoretical peptide libraries

Measuring the abundance of peptides using DIA approaches requires carefully crafted ion libraries. These libraries contain five key elements: the mass/charge (*m/z*) of the precursor ion (Q1), the *m/z* of the fragment ions (Q3), the RT of the precursor ion, the protein name, and the amino acid sequence of the precursor ion (Supplementary Table S1). Each Q1/Q3 pair (transition) at a specific RT should be unique and can be used for measurement of a specific peptide. Ion libraries for SWATH are typically constructed from protein identifications based on DDA data using database search software such as ProteinPilot. While this approach for generating ion libraries is efficient and convenient, these libraries will not contain peptides that were not identified by DDA analysis, such as low abundance peptides or those with PTMs not included in the search parameters. Therefore, libraries generated by this approach only include a subset of the potentially detectable peptides in a SWATH experiment. This limitation is especially severe for glycopeptides, because glycans are structurally heterogeneous and difficult to identify with standard proteomic database search methods [11, 12].

To overcome the limitation of DDA-driven libraries, tailored libraries can be constructed that include the desired Q1/Q3 pairs and corresponding RTs. Values for specific Q1/Q3 pairs and RTs can be obtained from manual exploration of the raw MS data, as we have previously described [5]. However, manual exploration of raw data and manual library construction is time consuming and error prone. Alternatively, libraries can be constructed with theoretical Q1/Q3 values. Here, we present DIALib, software that expedites and automates library construction for theoretical peptides and glycopeptides for use in DIA analysis (Figure 1).

**Figure 1.**
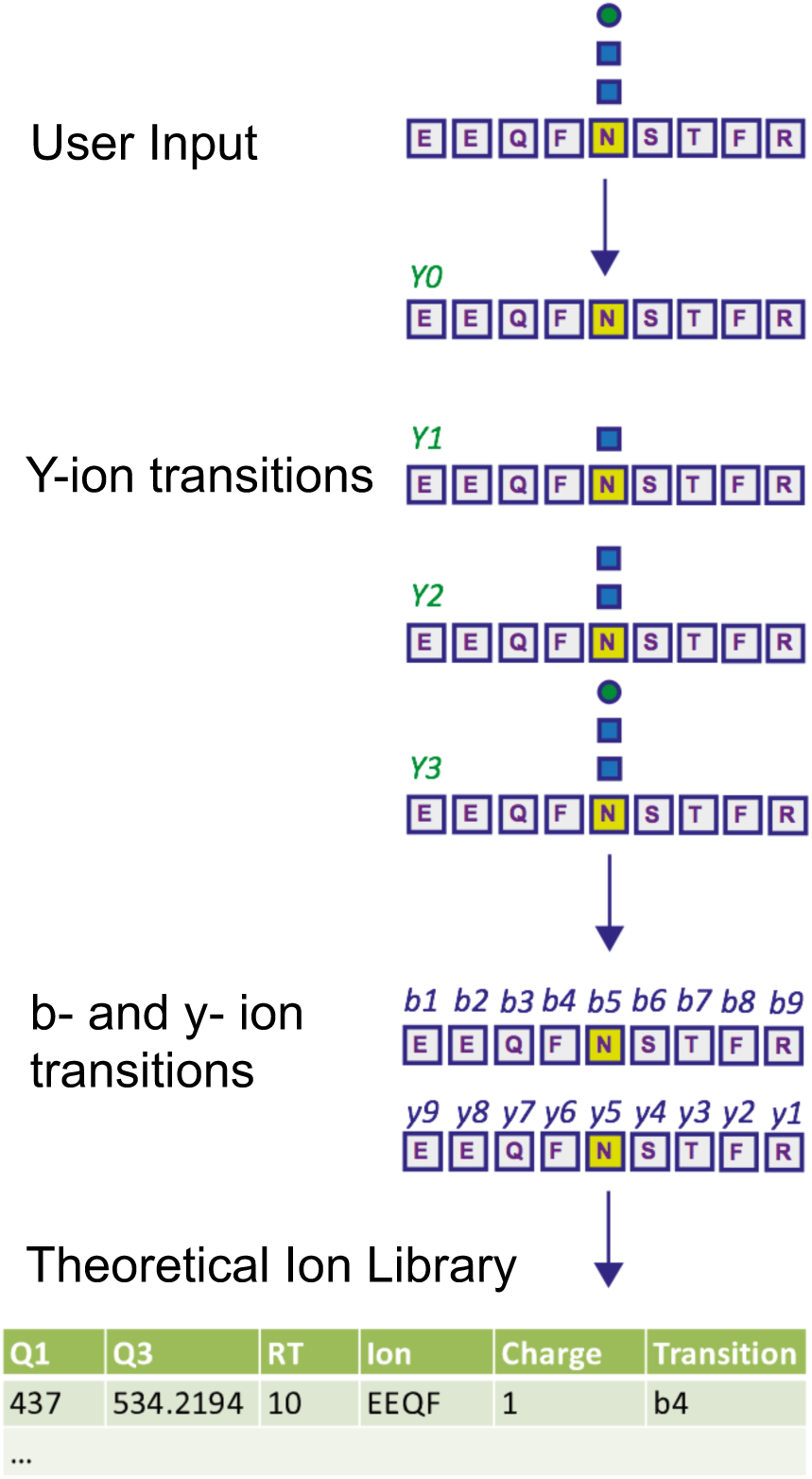
DIALib generates customized ion libraries for interrogation of post-translational modifications including protein glycosylation in Data Independent Acquisition MS data. The user inputs/selects the key parameters for construction of the ion library: amino acid sequence (or Uniprot ID), desired fixed and variable post-translational modifications, desired type of fragment ions (*b*, *y*, Y, or oxonium), retention time, and Q1. DIALib calculates the corresponding Q3 values and generates an ion library text file formatted for PeakView (SCIEX). If Y ions (e.g. HexNAc) are selected, DIALib searches for Asparagine (N) residues in the context of a sequon (e.g. NST, N in yellow) within the peptide sequences input, and constructs the Y series (Y0, no added sugars; Y1, with one HexNAc (blue square); Y2, with two HexNAc).

DIALib allows facile construction of SWATH ion libraries, including selection of pre-defined theoretical values and input of user-customized values for variables, including static modifications (alkylation of cysteine residues), non-glycan PTMs, Q1 (*m/z* windows), RT, *z*, and Q3 (*b*, *y*, Y, and/or oxonium ions) (Supplementary Table S1). One important feature of DIALib is the Q1 selection tool. During a SWATH experiment, the precursor *m/z* spectral space is divided into windows that can have the same or variable *m/z* size, typically with 1 *m/z* overlap between adjacent windows. DIALib allows the user to input arbitrary windows corresponding to any actual DIA experiment or to select a default fixed window method. Similarly, the user can input arbitrary customized RT values, or select one or more integer RT values from 1 to 60 min. With a focus on glycopeptide analysis, DIALib can construct libraries for any combination of *b*, *y*, Y, and oxonium ions, including libraries with only Y or oxonium ions. To generate a library with only Y ions, DIALib uses the peptide sequences provided by the user to calculate theoretical Y0 and Y1 (Hex(1)) Y ion transitions for *O*-man *O*-glycans, and Y0, Y1 (HexNAc(1)), and Y2 (HexNAc(2)) Y ion transitions for *N*-glycans and *O*-GalNAc *O*-glycans, considering *z* = 1 or 2 (up to a maximum of 6 Q3 values). Since all glycoforms (differently glycosylated versions of the same underlying peptide) share the same Y ions, DIALib generates a library that can potentially measure all possible glycoforms of the glycopeptides of choice in the entire spectral space sampled with SWATH [15]. A similar concept is applied when using only oxonium ions. We provide examples of both types of libraries below. DIALib can also be used to construct *by* libraries for glycopeptides, peptides with other modifications, or non-modified peptides.

### DIALib theoretical Y ion libraries can measure N-glycopeptides in a complex sample

We first tested if a Y ion library constructed with DIALib could reproduce results obtained with a manually curated glycopeptide library. To do this, we chose our previously published yeast cell wall glycoproteomics dataset (ProteomeXchange, PXD003091), in which we had measured site-specific glycan structure and occupancy in yeast cell wall glycoproteins using a manually curated ion library of *b*, *y*, and oxonium ions [5]. This moderately complex glycoproteome allowed clear measurement of glycopeptides with DDA and DIA approaches (Figure 2). We used DIALib to construct a Y ion library for the eight glycopeptides that were characterized in detail in this previous work [5]. We initially took a stringent theoretical approach, and made no assumptions about the *m/z* or RT of detectable versions of these glycopeptides. The Y ion library contained a maximum of 6 transitions (Y0, Y1, and Y2, and *z* = 1 or 2) for each of the 34 Q1s or *m/z* windows, and a fixed RT for all glycopeptides with a RT window covering the entire LC chromatogram (Supplementary Tables S1 and S3). Using the DIALib Y ion library, we re-interrogated the published SWATH dataset [5] and measured the abundance of each glycopeptide in each *m/z* window over the full LC elution timeframe. We exported the data allowing 100% FDR, to provide a complete data matrix (Supplementary Table S4). To facilitate comparison between yeast strains, we displayed the data as heatmaps (Figures 3-4, and Supplementary Figure S1). Figure 3 shows the results for four representative glycopeptides in the eight yeast strains. For each glycopeptide in Figure 3, two heatmaps are displayed. One heatmap shows the published data depicting the normalized summed *b*, *y*, and oxonium ion intensity for select precursors (Q1s) corresponding to several glycoforms per glycopeptide (in this case, glycoforms are glycopeptides carrying *N*-glycans with different number of mannose residues) (Figure 3B, E, H, and K) [5]. The second heatmap shows analyses from a DIALib library using normalized Y ion signal intensity for each *m/z* window corresponding to the published Q1s for each glycoform (Figure 3A, D, G, and J). To directly compare these libraries, we performed a correlation between each of the measurements from the DIALib Y ion and manual libraries shown in the heatmaps (Figure 3C, F, I, and L, and Supplementary Figure S1). The results of these analyses showed that for some glycopeptides, including Gas1 N^40^GS, Crh1 N^177^YT, and Gas1 N^95^TT, the output of both libraries was remarkably similar (R^2^ = 0.989, 0.968, and 0.912, respectively, Figure 3A-F and Supplementary Figure S1).

**Figure 2.**
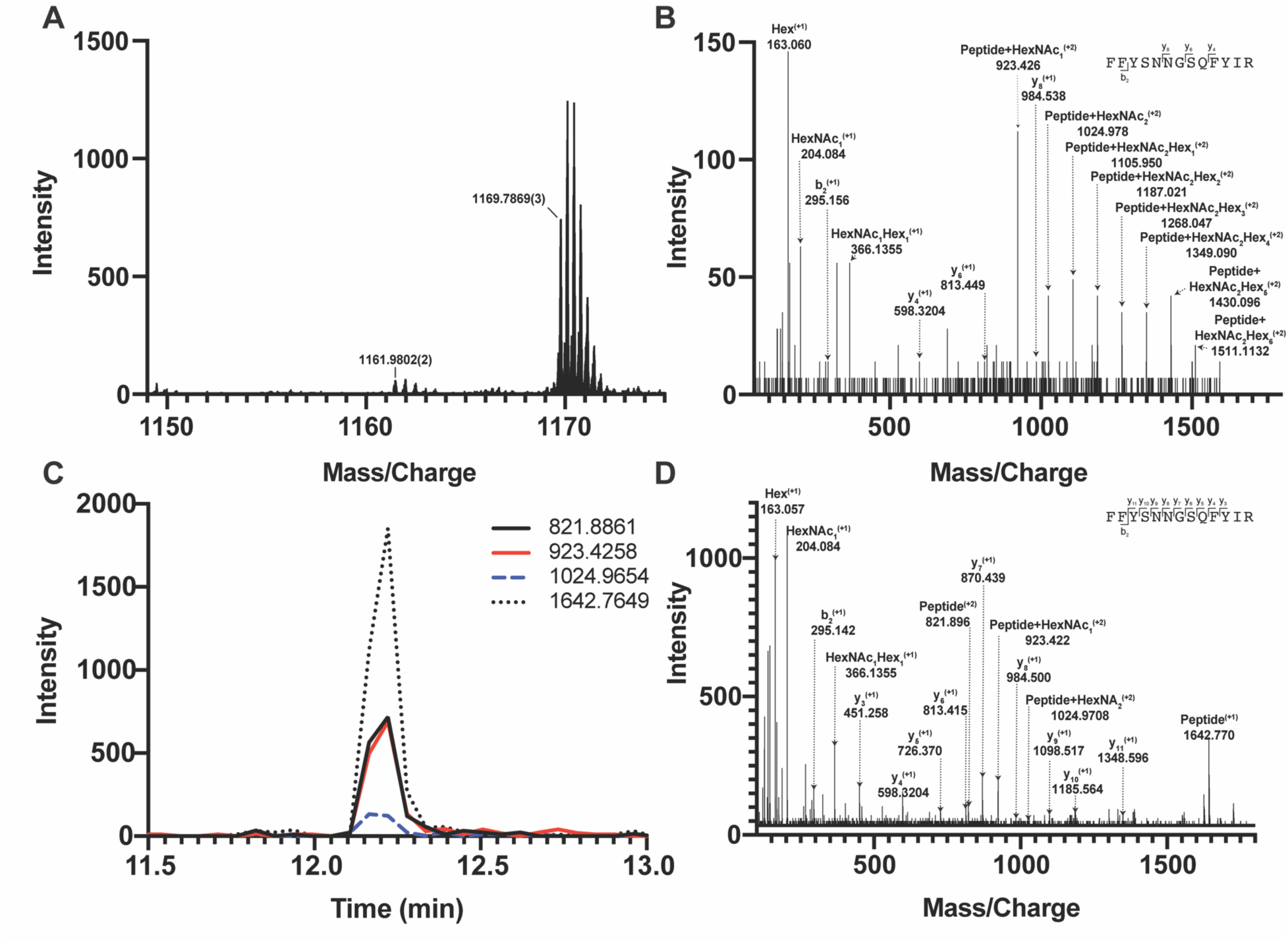
DDA and DIA detection and measurement of yeast glycopeptides. **(A**) DDA MS1 spectrum of a wild type cell wall sample at 12.195 min showing the precursors in the SWATH window from 1150-1175 *m/z*. Two main precursors fragment in this window: the Gas1N^40^GS glycopeptide at 1169.7869^3+^ *m/z* and a non-glycosylated peptide at 1161.9802^2+^ *m/z*. (**B**) DDA MS/MS spectrum of the Gas1N^40^GS glycopeptide at 1169.7869^3+^ *m/z* (Gas1 N^40^GS peptide with HexNAc2Man9). (**C**) DIA MS/MS extracted ion chromatograms for Gas1 N40GS transitions Y0 (821.8861^2+^ and 1642.7649^1+^), Y1 (923.258^2+^), and Y2 (1024.965^2+^) at 12.2 min in SWATH window 1149-1175 *m/z*. (**D**) DIA MS/MS spectra for the SWATH window 1149-1175 *m/z*. Fragment ions are labelled corresponding to transitions from the precursor 1169.7869^3+^ Gas1 N40GS peptide with HexNAc2Man9.

**Figure 3.**
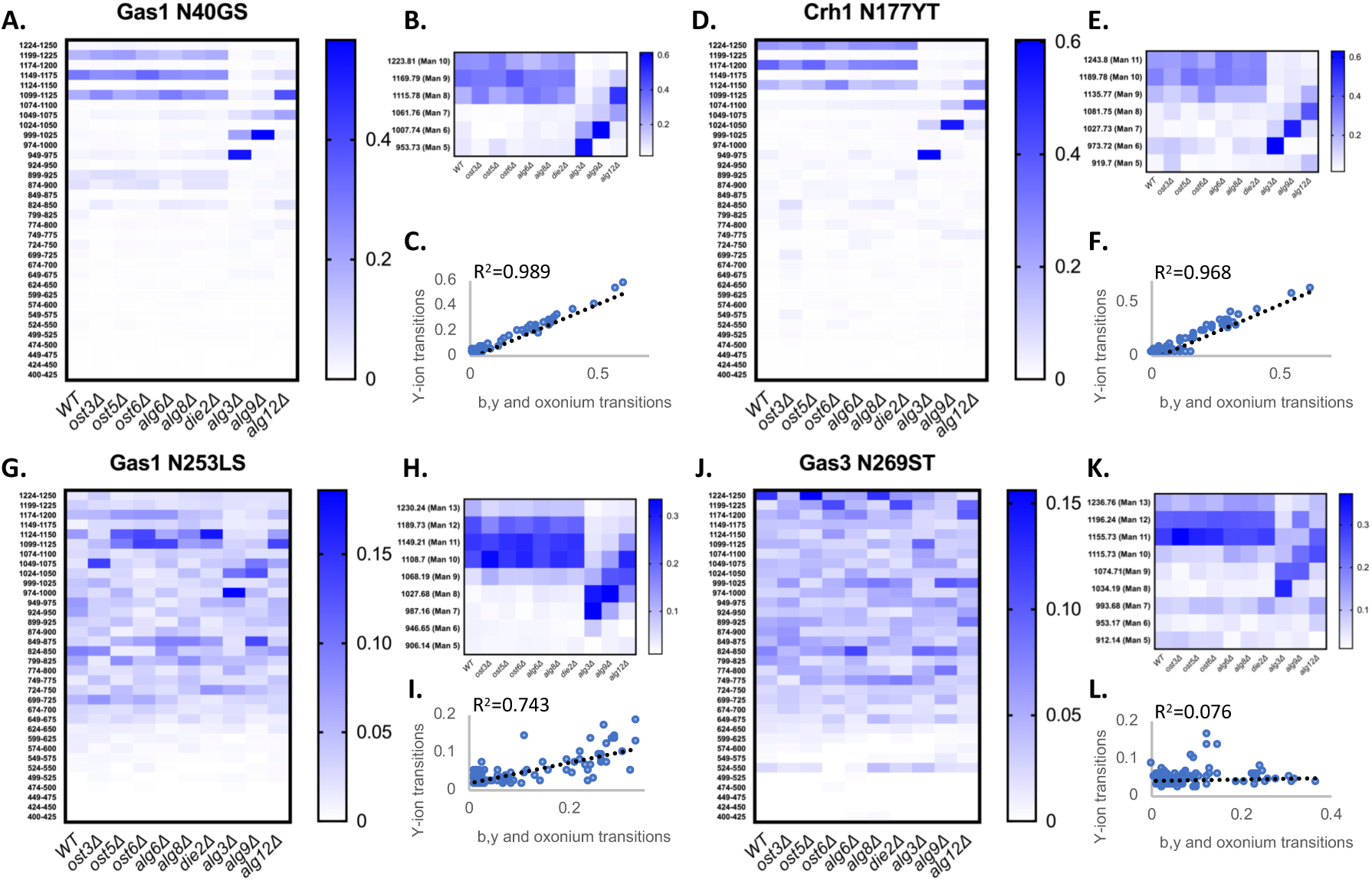
Detection and measurement of glycan structural heterogeneity in yeast glycopeptides is comparable using a DIALib Y ion library and a manually curated library. Abundance of glycoforms of glycopeptides containing Gas1 N^40^GS, Gas1 N^253^LS, Crh1 N^177^YT, and Gas3 N^269^ST measured using a DIALib Y-ion library (**A**, **D**, **G**, **J**) or a manually curated ion library [5] (**B**, **E**, **H**, **K**) in wild type yeast and yeast with mutations in the *N*-glycan biosynthesis pathway. Each square corresponds to the mean intensity of triplicate measurements. (**C**, **F**, **I**, **L**) Correlation between the data obtained using a DIALib Y ion library and a manually curated ion library [5], with linear fit and R^2^ values.

**Figure 4.**
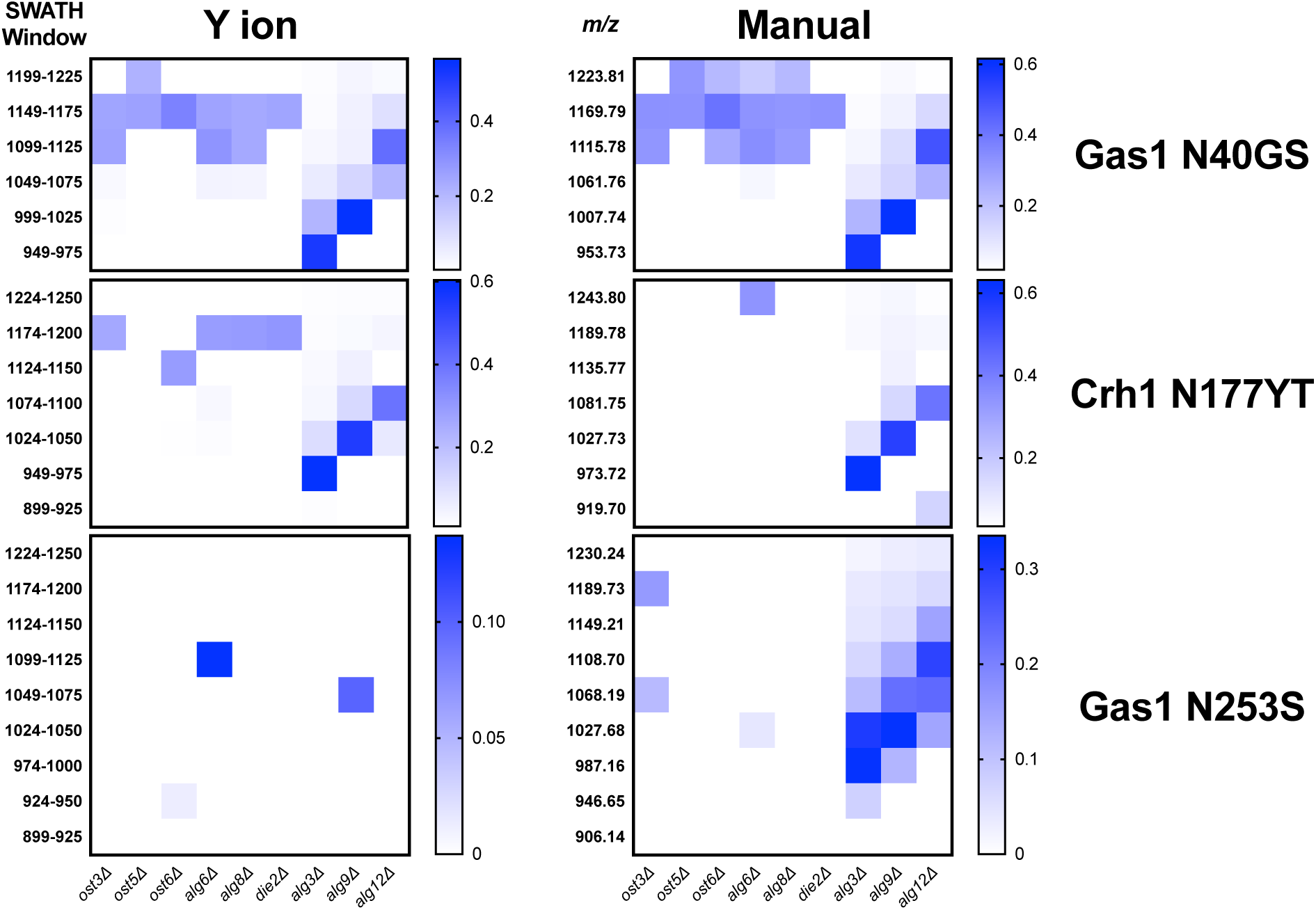
Theoretical DIALib Y ion library without experimental peptide retention times provides variably robust glycopeptide measurement. Abundance of glycoforms of glycopeptides containing Gas1 N^40^GS, Gas1 N^253^LS, and Crh1 N^177^YT measured using a DIALib Y ion library (left panels) or a manually curated library [5] (right panels) in wild type yeast and yeast with mutations in the *N*-glycan biosynthesis pathway. Each square corresponds to the mean intensity of triplicate measurements. Only statistically significant data is depicted, *p* < 0.05 when comparing glycoform abundance in any mutant strain to wild-type yeast.

Results for the other glycopeptides were variable. For example, the Y ion library was able to detect and measure many, but not all, of the glycoforms measured by the manual library for the glycopeptides Gas1 N^253^LS, Ecm33 N^304^FS, and Gas3 N^350^VS (R^2^ = 0.743, 0.665, and 0.427; Figure 3G-I, and Supplementary Figure S1). Interestingly, the glycoforms of Gas3 N^269^TS were detected and measured with the manual ion library but not by the Y ion library (R^2^ = 0.076, Figure 3J-L). Finally, glycopeptide Gas1 N^57^ET gave a poor glycoform profile with either the Y ion library or the published library (R^2^ = 0.084, Supplementary Figure S1). Thus, measurement of site-specific *N*-glycan structural heterogeneity in yeast cell wall glycoproteins using DIALib’s Y ion library were generally in agreement with our published data [5], with some inconsistencies. Together, these results indicated that DIALib’s Y ion library can be used to detect and measure *N*-glycopeptides in a moderately complex sample.

Robust measurement of peptides is critical in a DIA experiment. To compare the robustness of measurement of glycopeptide abundance using the Y ion library with that of the manually curated library [5], we focused on two glycopeptides that were similarly measured by the Y ion and manual libraries (Gas1 N^40^GS and Crh1 N^177^YT, R^2^ > 0.96, Figure 3A-F and Supplementary Figure S1), and one with an intermediate correlation score but showing a similar glycoform heatmap pattern to the manually curated library (Gas1 N^253^LS, R^2^ = 0.743, Figure 3G-I and Supplementary Figure S1). Figure 3 displays all the data, regardless of significance. We constructed a new set of heatmaps for the selected glycopeptides that only depicted data points that were statistically significantly different between wild type and mutant yeast strains (P < 0.05) (Figure 4). The measurements obtained with the manually curated library were robust, and identified many statistically significant differences in glycoform abundance between the wild type yeast and the glycosylation mutants, especially in the mannosyltransferase mutants (*alg3Δ*, *alg9Δ*, and *alg12Δ*) (Figure 4, Manual; note the similarity with the heatmaps in Figure 3B, E, and H) [5]. The measurements using the DIALib Y ion library showed mixed results (Figure 4, Y ion; compare to Figure 3A, 2D, and 2G). For example, the Gas1 N^40^GS and Crh1 N^177^YT glycopeptides (high R^2^, Figure 3A and 2D and Figure 4) displayed similar significant heatmap profiles to the manually curated library (compare Manual and Y ion heatmaps in Figure 4). On the other hand, the Gas1 N^253^LS glycopeptide (moderate R^2^, Figure 3G), displayed a different significant heatmap pattern to the one obtained with the manually curated library (Figure 3H and Figure 4), with almost no significant differences detected in the Y ion generated “significant” heatmap (compare Manual and Y ion in Figure 4, and compare with Figure 3G-I). Thus, for some glycopeptides, the Y ion measurements were robust, while for another glycopeptide the higher variability between samples led to identification of fewer statistically significant differences between the wild type and mutant yeast strains (Figure 4). Together, this suggested that in the absence of DDA data on peptide fragmentation and RT, DIALib’s Y ion library is an excellent tool to obtain an overview of the different glycoforms in a sample, but may not by itself be a robust method to search for statistically significant differences in glycoform abundance between complex samples.

### Enhancing glycoform measurement with DIALib b, y, and Y ion libraries

We wished to improve the ability of DIALib’s theoretical libraries to measure glycoforms in a complex sample. One difference between the published manual library and DIALib’s library was the number of transitions per glycopeptide. The manually curated libraries included 6 *b* and *y* transitions as well as the 204.08 (HexNAc) and 366.14 (HexNAc-Hex) oxonium ions for each glycopeptide [5]. In the Y ion library, the number of Y ion transitions is limited by the *m/z* of the peptide and by the mass selection range of the equipment. The DIALib libraries used in this work had up to a maximum of 6 Y transitions depending on the size of the underlying peptide (as described above). For four out of the five glycopeptides that showed an intermediate or low R^2^ in the DIALib *vs* manual library comparison (Gas1 N^57^ET, Gas3 N^269^ST, Gas1 N^253^LS, and Ecm33 N^304^FS, Figure 3G-I and Supplementary Figure S1) the DIALib Y ion library contained only three transitions. The remaining three glycopeptides with high R^2^ had four or more Y ion transitions per glycopeptide (Gas1 N^40^GS, Crh1 N^177^YT, and Gas1 N^95^TT, Figure 3A-F and Supplementary Figure S1). We hypothesized that incorporating additional transitions to the Y ion library would improve glycopeptide detection and measurement. To test this idea, we used DIALib to generate four additional ion libraries for the same eight glycopeptides as above: 1) a *by* ion library containing all theoretical *b* and *y* ions for the nonglycosylated version of each glycopeptide (excluding *b1*, *y1*, and MH-H_2_O ions and including the MH ion); 2) a stop*by* ion library containing all theoretical *b* and *y* ions for each glycopeptide that did not include the glycosylated asparagine residue (except *b1*, *y1*, and MH-H_2_O ions); 3) a *by*+Y ion library which combined the *by* and Y ion libraries; and 4) stop *by*+Y ion library which combined the stop*by* and Y ion libraries (Supplementary Table S5). Excluding fragment ions containing the asparagine residue in the stop*by* libraries ensured that only transitions common to the unglycosylated and glycosylated forms of the same peptide were present in the library.

Following the procedure described above for the Y ion library, we constructed the four new DIALib libraries for the eight glycopeptides, measured the abundance of each glycopeptide in each *m/z* window over the full LC elution timeframe, and exported the data allowing 100% FDR. Surprisingly, the results showed that none of the four new libraries outperformed the Y ion library or the manually curated library for measurement of glycoform abundance in a complex sample (Supplementary Figure S2). In addition, we observed that the combination of the *by* and stop*by* transitions with the Y ion was generally detrimental to measuring glycoform abundance compared to the *by*, stop*by*, and Y ion libraries alone. The only notable exception was for the Crh1 N^177^YT glycopeptide, for which the incorporation of the Y ions helped in glycoform measurement compared to the *by* and stop*by* libraries alone (Supplementary Figure S2). Of the four libraries, the stop*by* was more successful in producing a glycoform heatmap pattern resembling the published one (compare Supplementary Figures S1 and S2). To note, one key feature of the *by* and stop*by* libraries (not shared by the Y ion library) was that they had the potential to also allow measurement of the unglycosylated peptide, providing simultaneous measurement of glycan occupancy and structure. This was especially evident in the stop*by* library, in which the unglycosylated peptides were identified for 6/8 of the glycopeptides measured (Crh1 N^177^YT, *m/z* window 512; Ecm33 N^304^FS, *m/z* window 787; Gas1 N^40^GS, *m/z* windows 812 and 537; Gas1 N^95^TT, *m/z* window 587; Gas1 N^57^ET, *m/z* window 1031, and Gas3 N^350^VS, *m/z* window 437, Supplementary Figures S2 and [5]).

The poor performance in glycoform measurement of the *by/*stop*by* libraries compared to the Y ion libraries was unexpected. We considered two possibilities that might explain the inaccurate peak picking that was the underlying cause of this poor performance: that the *by/*stop*by* libraries used were not analytically optimal, as they contained all (or most) theoretical *by/*stop*by* fragment ions for each glycopeptide; or that the RT window used was too wide, as it covered the entire LC gradient. Indeed, we observed that reducing the number of transitions in the *by* library while maintaining a large XIC RT window tended to enhance peak peaking (data not shown). However, the optimal number of transitions required for optimal peak picking was peptide-dependent (data not shown), and thus could not be set as a fixed parameter for all glycopeptides. Next, we tested if narrowing the XIC RT window would improve the performance of the *by,* stop*by,* stop*by*+Y, and Y libraries, while maintaining all theoretical transitions in the library. Since a narrow XIC RT window requires an accurate RT for each peptide, we constructed *by,* stop*by,* stop*by*+Y, and Y libraries using the manually optimized RT from the published library [5] (Supplementary Tables S6 and S7). We re-analyzed the data with these RT-optimized libraries using an RT XIC window of 2 min (Figure 5 and Supplementary Figures S3). The results showed that when the search space was constrained due to a more specific RT, a reduced XIC RT window, or lower number of transitions, the measurement of peptides improved (Figure 5 and Supplementary Figures S3). First, all the new RT-optimized libraries were able to successfully identify and measure glycopeptides that were not measured with the non RT-optimized libraries (compare Supplementary Figure S2 with Figure 5 and Supplementary Figure S3). This improved peak picking performance was especially evident for the Gas3 N^269^ST glycopeptide, since the stop*by*, stop*by*+Y, and Y RT-optimized libraries successfully measured glycoforms for Gas3 N^269^ST, while all the non RT-optimized libraries failed to do so (compare Figures 3 and 5). In general, the heatmaps obtained using the stop*by* library had lower background than when the *by* library was used (i.e. no measurements where no peptides were expected), especially when an optimized RT was employed (Figure 5 and Supplementary Figures S2 and S3). The improvement in peak picking for the RT-optimized libraries was also demonstrated by the higher R^2^ correlation scores obtained when comparing site-specific glycan structural heterogeneity between the RT-optimized libraries and the manually curated library (Figure 5B). Of all the libraries used in this study, the RT-optimized stop*by*+Y library was the best performing, not only because it consistently displayed one of the highest overall R^2^ correlation scores for all 8 glycopeptides (0.84 +/− 0.14, Figure 5C), but also because it could detect and measure both glycan occupancy and structure simultaneously (Figures 3-5 and Supplementary Figures S1 and S2). Nevertheless, it is important to point out that in the absence of prior RT information, the Y ion library still considerably outperformed all other libraries tested in this work (Figure 3 and Supplementary Figure S2). Overall, these results indicated that in order for the DIALib libraries to optimally detect and measure glycoforms in a complex sample, prior optimization of the RT for each peptide and a narrow XIC window are required.

**Figure 5.**
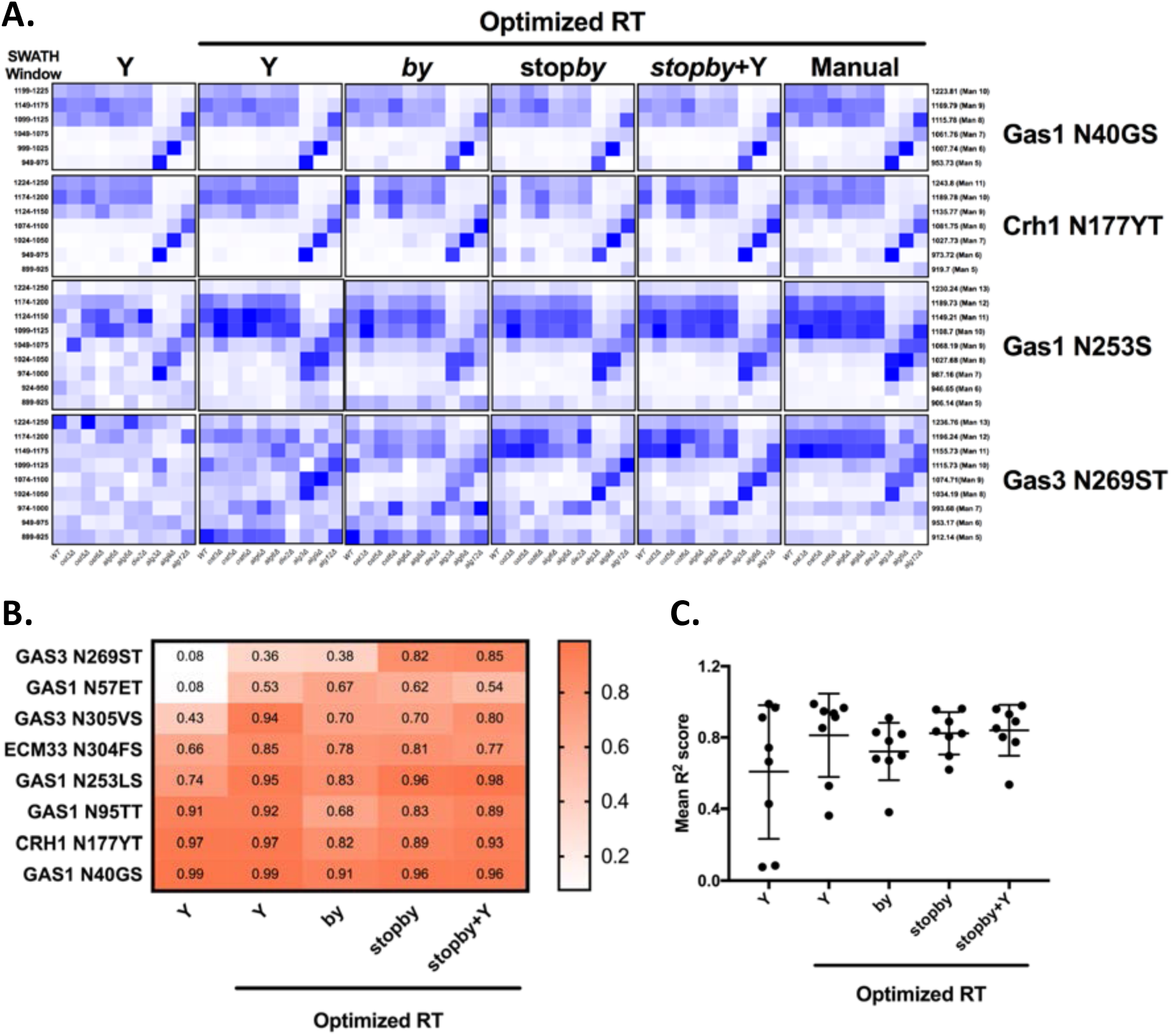
Optimized retention times improve peak picking for DIALib libraries. (**A**) Abundance of glycoforms of glycopeptides containing Gas1 N^40^GS, Crh1 N1^77^YT, Gas1 N^253^LS, and Gas3 N^269^ST measured using the *by*, stop*by*, stop*by*+Y, and Y DIALib libraries and manually curated library incorporating experimental peptide retention times [5], measured in wild type yeast and in yeast with mutations in the *N*-glycan biosynthesis pathway. Each square corresponds to the mean intensity of triplicate measurements. Only windows corresponding to the expected *m/z* for the different glycoforms are depicted. (**B**) Heatmap and (**C**) plot of R^2^ values for correlations of the intensity of each glycoform in all yeast strains, comparing each DIALib library to the manually curated library [5].

### DIALib Y ion libraries detect mammalian N-glycopeptides

To test the ability of the DIALib Y ion libraries to identify glycoforms in a sample of mammalian glycopeptides, we prepared tryptic digests of purified human serum Immunoglobulin G. The Y ion library was constructed for *N*-linked sequon-containing tryptic glycopeptides from IgG1 (EEQYN^180^STYR), IgG2 (EEQFN^176^STFR), IgG3 (EEQYN^227^STFR), and IgG4 (EEQFN^177^STYR). The sequon containing peptides for IgG3 and IgG4 are isobaric and cannot be distinguished using a Y ion library. We constructed the DIALib Y ion library to span the entire DIA space, searching all 34 *m/z* windows with 1 min XIC RT windows across the full LC timeframe (Supplementary Table S8). This approach also allowed measurement of glycoforms with *m/z* values that fell in the same *m/z* window, but whose glycan structures altered the glycopeptide RT. To process the data, we first normalized the signal intensities of each Q1/RT pair to the total signal intensity for the sample (all Q1 at all RT). We organized the normalized data as a RT *vs m/z* window matrix (Figure 6). To simplify the matrix due to differences in the observed RT at different Q1s, we rounded up the RTs for each measurement to the nearest integer (using an in house designed script reformat_peptide3.py) (Figure 6 and Supplementary information). Figure 6 shows heatmaps summarizing the results for the IgG glycopeptides. The majority of the signal for IgG1 was concentrated at RT 7 min in multiple Q1s, suggesting that multiple glycoforms of the same IgG1 glycopeptide eluted at 7 min. Similarly, most glycoforms of IgG2 eluted at 8 min (Figure 6). As expected, the Y ion glycan profile for IgG3 and IgG4 was identical because their glycopeptides have isobaric sequences (Figure 6). We validated this DIALib derived data by identifying IgG glycopeptides using Byonic in the DDA LC-MS/MS data (Supplementary Tables S2 and S9) and by manual inspection of the raw data. The manually validated Byonic glycopeptide identifications generally aligned well with the data obtained with the Y ion library. For IgG1 and IgG2 glycopeptides, there was clear signal detected in almost all *m/z* windows for which glycopeptides were identified by Byonic (Supplementary Table S2 and S9). Byonic searches did not identify any glycoform for the EEQYNSTFR peptide from IgG3/4, possibly because of the lower abundance of IgG3/4 compared to IgG1 and IgG2 (Supplementary Table S9). These results highlight the potential improved sensitivity of DIA compared with DDA analyses. Together, the data demonstrated that the DIALib Y ion library can successfully profile, detect, and measure mammalian *N*-glycopeptides.

**Figure 6.**
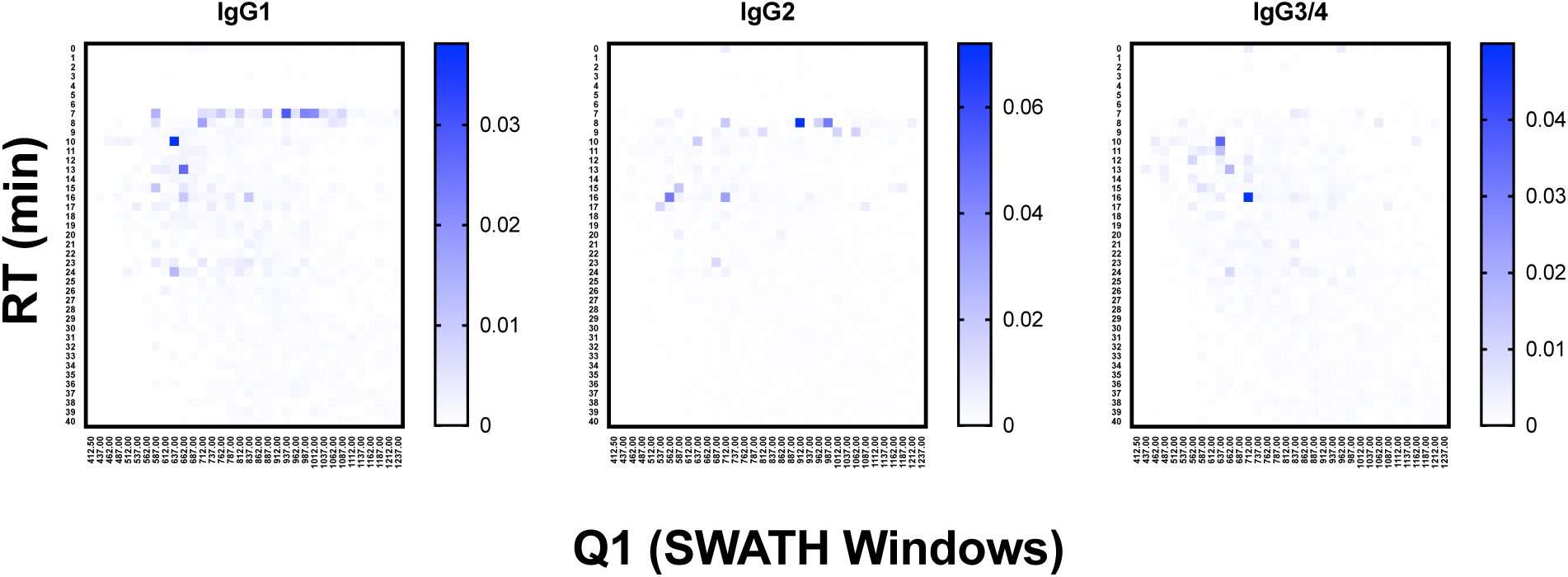
Glycoprofiling human Immunoglobulin G using a DIALib Y ion library. Glycopeptide profiles across all retention times and *m/z* windows for each IgG isotype.

### DIALib libraries for O-glycopeptides

DIALib can also generate theoretical Y ion libraries to profile the site-specific structural diversity of the *O*-glycoproteome. ProteinPilot and Byonic searches of our published yeast cell wall dataset [5] identified multiple glycoforms of *O*-glycopeptides (Supplementary Tables S10 and S11, respectively). We constructed a DIALib library focusing on one *O*-glycopeptide: T^318^SGTLEMNLK^327^ from Hsp150 (Supplementary Table S12), that is predicted to be *O*-glycosylated at positions T^318^, S^319^, and/or T^321^ [19] (Supplementary Tables S10 and S11). Similar to the Y ion library for *N*-glycans described above (Supplementary Tables), the *O*-glycopeptide DIALib library included Y0 and Y1 transitions (assuming only one modified site) for T^318^SGTLEMNLK^327^ for 34 Q1s and all RT. A 2 min XIC RT window over the full LC was used to identify and measure the *O*-glycoforms (Supplementary Table S13). As described above for the IgG analysis, we arranged the normalized data into an RT *vs m/z* window matrix for each yeast strain. Figure 7 shows an example of the RT *vs* Q1 matrix for the WT strain, although the heatmaps for all the strains were very similar (Figure 7, top panel). The four glycoforms of the Hsp150 *O*-glycopeptide (Man(1) to Man(4)) identified by Byonic as eluting at an RT ∼10 min (Supplementary Table S11) were also measured in all strains using the DIALib library (Figure 7, bottom panel). We manually confirmed the presence of the corresponding Y and Hex oxonium ions (145.05 and 163.06) for these glycoforms in the *m/z* windows 637 (Man(1)), 712 (Man(2)), 787 (Man(3)), and 862 (Man(4)) (Figure 7). We also observed that the Man(2) glycoform of the T^318^SGTLEMNLK^327^ peptide was the most abundant of the glycoforms (Figure 7, *m/z* window 699-725). Therefore, DIALib can also detect and measure *O*-glycopeptides in a moderately complex sample.

**Figure 7.**
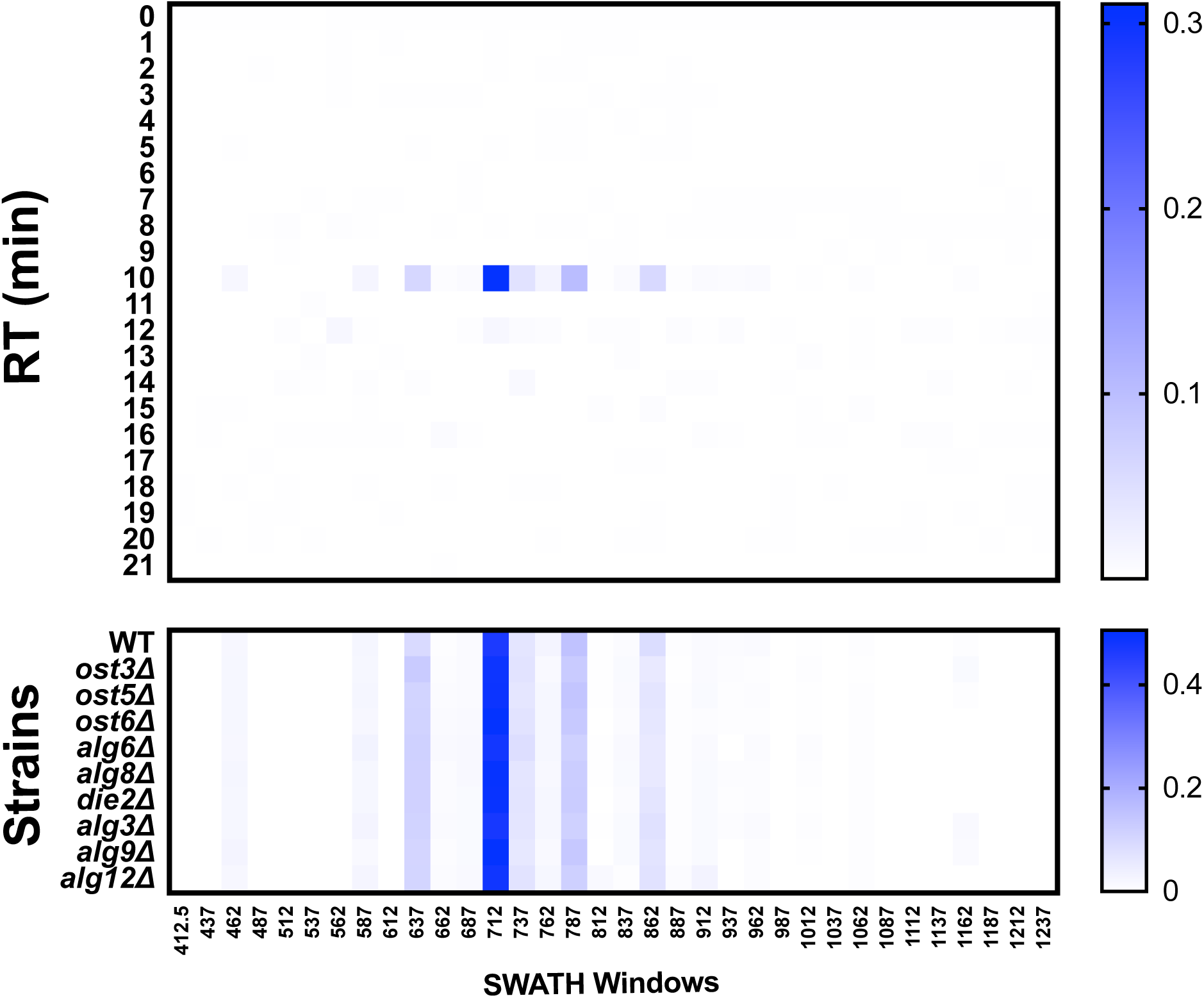
*O*-glycoprofiling yeast cell wall glycopeptides. (**A**) Glycopeptide profile across all retention times and *m/z* windows with a DIALib Y ion library for the HSP150 *O*-glycopeptide T^318^SGTLEMNLK in wild type yeast. The mean (n=3) normalized signal intensity at each *m/z* window and each retention time is shown for all *m/z* windows in the entire LC gradient. (**B**) Glycopeptide profile of the Hsp150 *O*-glycopeptide T^318^SGTLEMNLK at a retention time of 10 min for wild type yeast and glycosylation mutants.

### Profiling the “oxoniome”

During glycopeptide fragmentation in CID/HCD mode, glycans fragment into smaller ions called oxonium ions [20]. Oxonium ions not only signal the presence of a glycan, but can also provide some information on glycan composition and structure [21]. Thus, measurement of oxonium ion abundance can be used to detect genetic or environmental conditions that impact protein glycosylation. Comprehensive profiling of the “oxoniome” of a sample can be performed with no *a priori* knowledge of the sample’s contents by measuring oxonium ion abundance across the entire *m/z* and RT range. We used DIALib to create a library that would measure the eight most common Hex and HexNAc oxonium ions (Q3) at every minute in all *m/z* windows. The oxonium ions included were: Hex-(H_2_O)_2_ (127.0389 *m/z*), HexNAc-(H_2_O)_2_-COH_2_ (138.055 *m/z*), Hex-H_2_O (145.0495 *m/z*), Hex (163.0601 *m/z*), HexNAc-(H_2_O)_2_ (168.0655 *m/z*), HexNAc-H_2_O (186.0761 *m/z*), HexNAc (204.0866 *m/z*), and HexNAc-Hex (366.1395 *m/z*) (Supplementary Table S14). DIALib includes a standard catalogue with additional oxonium ions such as from sialic acids, which we did not include in this analysis of the yeast cell wall glycoproteome. We used this library to profile cell wall samples from wild type yeast and the nine glycosylation mutants. As above, we searched the entire LC gradient and *m/z* window range using an XIC RT window of 2 min (Supplementary Table S15). To profile the overall oxoniome, we summed the signal intensity of all eight oxonium ions in each RT and *m/z* window (30 retention times x 34 *m/z* windows = 1020 data points per strain) and then normalized this value to the total oxonium signal for each strain (sum of the 1020 data points per strain). The data was rearranged to depict a heatmap of RT *vs m/z* windows (Figure 8). Figure 8B shows the results of oxonium ion profiling as heatmaps for wild type yeast, and the *ost3Δ* and *alg3Δ* mutants. Most of the signal detected was concentrated in high *m/z* windows between 5-20 min, as expected for typical *m/z* and elution characteristics of glycopeptides (Figure 8B). Ost3 is a subunit of the OTase, and the *ost3*Δ mutant therefore transfers the same glycan to proteins as wild type, but at lower efficiency [5]. This is reflected in the relatively minor differences in oxoniome profile between wild type and *ost3*Δ cells. A useful way to depict differences between strains is the use of substraction heatmaps, in which each data point is the normalised intensity measured for wild-type less the normalised intensity measured for a given strain (Figure 8C). In agreement with the data obtained using the site-specific glycopeptide ion libraries above (Figures 1-4) and [5], we observed that mutant strains that had small effects on glycosylation occupancy and glycan structure also displayed small differences to wild type yeast in their oxonium substraction maps (e.g. the *die2Δ* and *ost5Δ* strains, Figure 8C). Also as before, we observed the mannosyltransferase mutants *alg3Δ*, *alg9Δ*, and *alg12Δ*, and the OST mutant *ost3Δ* displayed the highest difference compared with wild type (Figure 8C). We could also use oxonium ion data to profile the overall monosaccharide composition of the glycoproteome in each yeast strain. Consistent with the expected high mannose *N*-glycans and *O*-glycans in yeast glycoproteins, we measured higher signal for Hex than for HexNAc oxonium ions (Figure 8D). Strains lacking the mannosyltransferases Alg3, Alg9, and Alg12 transfer truncated *N*-glycan to protein (GlcNAc(2)Man(5), GlcNAc(2)Man(6), or GlcNAc(2)Man(8), respectively) (Figures 1-4) and [5]. We therefore predicted that oxoniom profiling would detect a different ratio of Hex:HexNAc in these strains compared to wild type yeast. Indeed, we observed statistically significant differences in the proportion of the corresponding oxonium ions between wild type and the *alg3Δ* and *alg9Δ* strains (Figure 8D). In these strains, the relative abundance of Hex oxonium ions was significantly lower and of HexNAc oxonium ions significantly higher than in wild type (P < 0.05, Figure 8D). Together, profiling the oxoniome with DIALib libraries was able to detect the expected compositional differences in the glycoproteomes of yeast glycan biosynthesis mutants, demonstrating that this approach provides a rapid and robust method of profiling glycosylation differences between samples.

**Figure 8.**
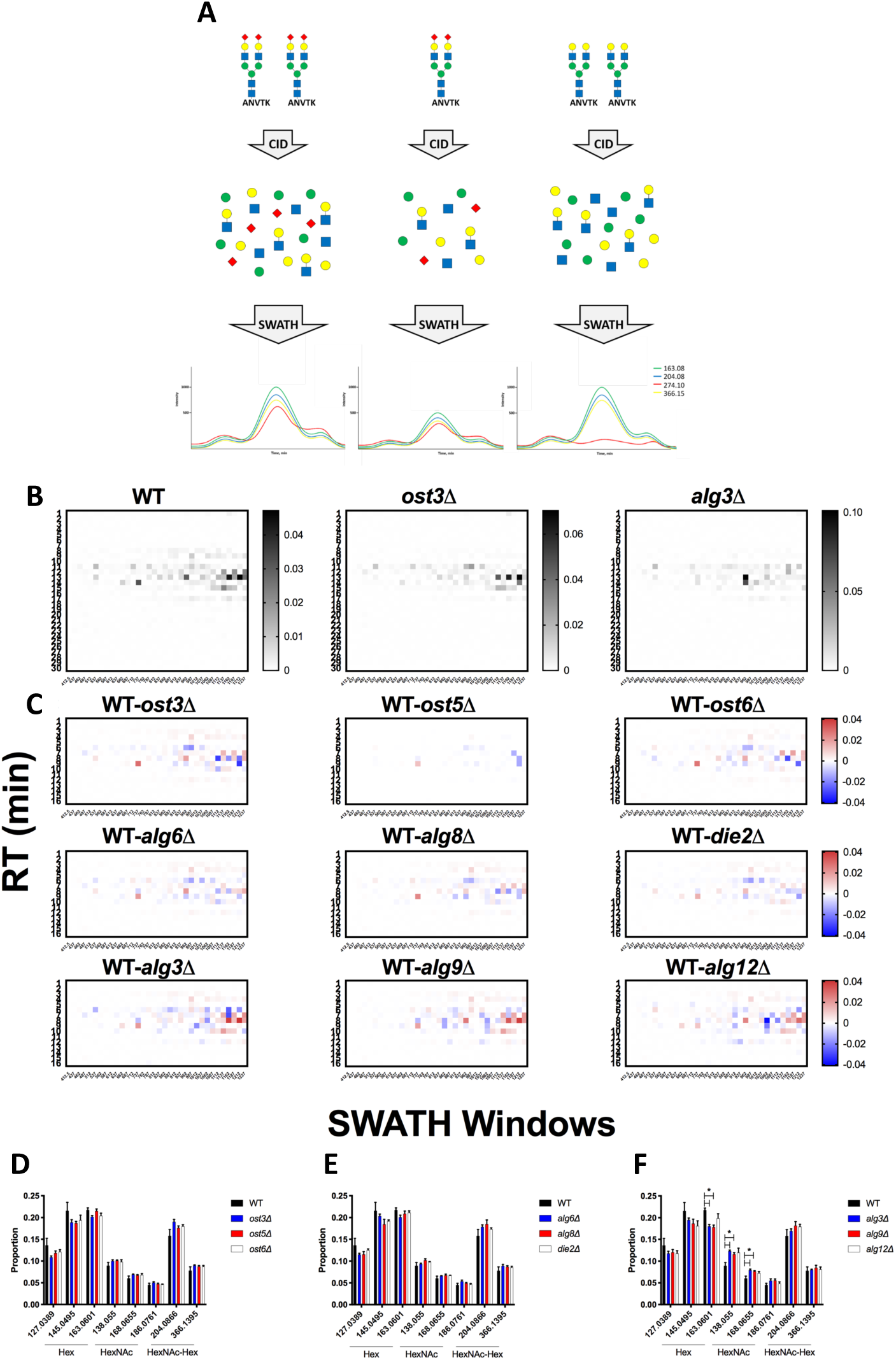
Oxoniome profiling of glycopeptides. (**A**) Cartoon showing how measuring oxonium ions with DIA can provide information on glycan monosaccharide composition and relative abundance. (**B**) Hex and HexNAc abundance profile in glycopeptides across all retention times and *m/z* windows with a DIALib oxonium ion library in glycopeptides from cell wall proteins of wild type and *alg3Δ* and *ost3Δ* glycosylation mutant yeast [5]. (**C**) Subtraction oxonium ion profiles for each glycosylation mutant strains compared to wild type. Relative abundance of Hex and HexNAc oxonium ions across all retention times and *m/z* windows for wild type and yeast with mutations in (**D**) oligosaccharyltransferase subunits, (**E**) glucosyltransferases, and (**F**) mannosyltransferases. Mean +/− SEM of triplicates. * indicates P < 0.05.

## Discussion

At its core, a DIA ion library is a collection of MS/MS profiles of peptides of interest. A high quality library should be able to replicate key features of the actual MS/MS profile of a peptide of interest, and most DIA libraries are therefore based on experimental DDA peptide identification. However, peptides with structurally complex post-translational modifications are often difficult to identify with standard DDA workflows. DIALib, as described, offers an alternative approach for building DIA libraries based on theoretical MS/MS fragmentations of the peptides with potentially complete coverage regardless of abundance.

Post-translational modifications very often alter the mass of the underlying peptide, while keeping much of the fragmentation profile of the peptide constant. This allows measurement of differently modified versions of the same underlying peptide using consistent MS/MS fragment ions, but at variable parent ion *m/z* values. DIALib can take advantage of this feature of modified peptides by generating a library that can measure peptides of interest across the full range of *m/z* windows. Peptide ions are often present at multiple charge states, and these can also be detectable using DIALib libraries, although care must be taken to distinguish peptide forms resulting from different charge states and those resulting from modifications. To achieve complete coverage of the space in a DIA experiment, DIALib can also provide a library to measure peptides of interest throughout the entire LC chromatogram by repeating each peptide entry with multiple retention times. To enable this unusual workflow, we labelled peptide sequences with “modification tags” that represent different combinations of retention time and *m/z* window. This approach allowed us to detect and measure glycoproteomic features missing from DDA-based analyses, including low abundance glycopeptides, and the “oxoniome” profile of a sample.

In using DIALib to aid analysis of test samples including a moderately complex yeast cell wall glycoproteome and purified human glycoproteins, we identified several strengths and limitations of the approach. DIALib’s completely theoretically generated ion libraries could be used to detect and measure *N*-glycopeptides in a moderately complex sample of yeast high mannose glycoproteins, although analytical performance was improved by incorporating DDA data on peptide fragmentation and RT. Accurate knowledge of peptide RT was an important parameter, suggesting that incorporation of tools to predict glyco/peptide elution behaviour would be a valuable addition to the approach. Analysis of *N*-glycopeptides from a simple mixture of mammalian glycoproteins showed that DIA glycopeptie analysis can have higher sensitivity than DDA, and that a theoretical DIALib ion library could successfully profile, detect, and measure mammalian *N*-glycopeptides. We also demonstrated that DIALib can detect and measure yeast mannose-rich *O*-glycopeptides in a moderately complex cell wall fraction. We note that site-specific analysis of more complex glycoproteomes may be more challenging than the examples presented here, especially for glycoproteomes with mammalian glycans with complex and heterogeneous structures. Care should also be taken in using DIALib with large numbers of predicted glycopeptides, as it may be difficult to effectively control false discovery rates. Finally, our approach for using DIALib libraries to profile the global oxoniome of yeast glycan biosynthesis mutants was able to detect the expected differences in their glycoproteomes. This global oxoniome profiling approach therefore provides a rapid and robust method of profiling glycosylation composition from all detectable analytes.

Automatically generated DIA ion libraries, either theoretical or sourced from experimental DDA data, are robust and reproducible. However, manual optimization of libraries may sometimes be necessary. In the case of a DIALib theoretical ion library, it is possible to further customise the library to increase sensitivity, specificity, or general analytical robustness, such as by manually selecting fragment ions for inclusion/exclusion, or including information on fragment intensity. However, such optimisation is peptide- and sample-specific, and requires expertise and careful manual implementation.

The analyses of glycoproteomes measured by DIA using libraries generated by DIALib that we present here clearly shows that this approach can be used to detect and measure site-specific modifications and profile the global glycoproteome. To develop this approach we used moderately complex yeast glycoproteome samples and purified human IgG glycoproteins. Measurement of site-specific modifications will require more care in glycoproteomes with higher complexity, such as complex human glycoproteomes with multiply sialylated and/or fucosylated complex N-glycan structures, while global oxoniome compositional glycoproteome profiling will remain a powerful approach even in complex glycoproteomes. We anticipate that DIALib will enable DIA glycoproteomics as a highly complementary analytical approach to DDA glycoproteomic strategies.

## Supporting information

Supplementary Information script reformat_peptide3.py

## Acknowledgements

We thank Dr Cassandra Pegg for preparing the IgG samples. We thank Dr Amanda Nouwens and Peter Josh at The University of Queensland, School of Chemistry and Molecular Biosciences Mass Spectrometry Facility for their assistance and expertise.

## Supplementary Material

**Supplementary Script S1. reformat_peptide3.py.** Round up RT and rearrange the datatable into RT vs windows matrix.

### Supplementary Tables

**Supplementary Table S1.**
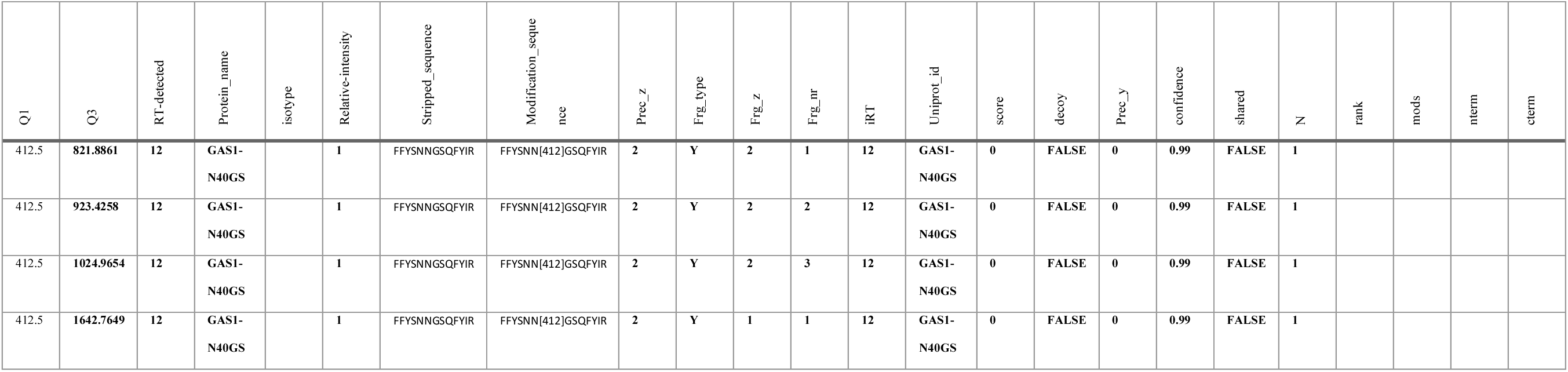
Example of details from a DIALib Y ion library formatted for use in Peakview. One Q1 value (out of 34) is depicted as an example, corresponding to the *m/z* window 400-425 *m/z* (average = 412.5 *m/z*). This example includes four theoretical Y ions (Q3s) calculated for the Gas1 N^40^GS peptide FFYSNNGSQFYIR (from a maximum of 6). The protein_name and the Uniprot_ID have the same identifier. The isotype column is left blank. Stripped sequence is the amino acid sequence of the peptide without modifications. The Modification_sequence is the amino acid sequence incorporating a modification in [], in this case it contains the Q1. Prec_z and Frg_z indicate the charge state of the precursor and fragment ions, respectively. Prec_z is arbitrarily always set as 2. Frg_type can be *b*, *y*, and Y. Frg_nr is the fragment ion number. The values for RT_detected and iRT are the same, and in this example they are set to 12 min. Score is set to 0, decoy as FALSE, Prec_y as 0, confidence as 0.99, Shared as FALSE, N as 1; and the remaining columns are left blank.

**Supplementary Table S2.** Select IgG glycopeptides identified by Byonic and detected by DIA using a DIALib Y ion library.

| Byonic |  |  |  |  |  |  | Y ion DIALib |  |  |  |  |
| --- | --- | --- | --- | --- | --- | --- | --- | --- | --- | --- | --- |
| Glycans<br>NHFAGNa | Glycan<br>type | Modification<br>Type(s) | Observed<br>m/z | z | Protein Name | Scan<br>Time | Peptide | Precursor<br>MZ | Precursor<br>Charge | Observed<br>RT | Normalized<br>intensity |
| HexNAc(4)Hex(4)Fuc(1) | G1F | N[+1607] | 932.696 | 3 | >sp\|P01857\|IGHG1 | 7.1108 | EEQYN[7.937]STYR | 937 | 2 | 7.1428 | 0.0294 |
| HexNAc(5)Hex(3)Fuc(1) | G0FN/B | N[+1648] | 946.371 | 3 | >sp\|P01857\|IGHG1 | 7.1453 | EEQYN[7.937]STYR | 937 | 2 | 7.1428 | 0.0294 |
| HexNAc(4)Hex(5)Fuc(1) | G2F | N[+1769] | 986.718 | 3 | >sp\|P01857\|IGHG1 | 7.094 | EEQYN[7.987]STYR | 987 | 2 | 7.1552 | 0.0222 |
| HexNAc(5)Hex(4)Fuc(1) | G1FN/B | N[+1810] | 1000.388 | 3 | >sp\|P01857\|IGHG1 | 7.1511 | EEQYN[7.1012]STYR | 1012 | 2 | 7.1524 | 0.0208 |
| HexNAc(4)Hex(5)Fuc(1)NeuAc(1) | G2FS | N[+2060] | 1083.742 | 3 | >sp\|P01857\|IGHG1 | 7.2462 | EEQYN[7.1087]STYR | 1087 | 2 | 7.2472 | 0.0070 |
| HexNAc(4)Hex(3)Fuc(1) | G0F | N[+1445] | 868.017 | 3 | >sp\|P01859\|IGHG2 | 8.3287 | EEQFN[8.862]STFR | 862 | 2 | ND | ND |
| HexNAc(4)Hex(4)Fuc(1) | G1F | N[+1607] | 922.041 | 3 | >sp\|P01859\|IGHG2 | 8.3881 | EEQFN[8.912]STFR | 912 | 2 | 8.3572 | 0.0720 |
| HexNAc(5)Hex(3)Fuc(1) | G0FN/B | N[+1648] | 935.708 | 3 | >sp\|P01859\|IGHG2 | 8.4446 | EEQFN[8.937]STFR | 937 | 2 | ND | ND |
| HexNAc(4)Hex(5)Fuc(1) | G2F | N[+1769] | 976.053 | 3 | >sp\|P01859\|IGHG2 | 8.3563 | EEQFN[8.987]STFR | 987 | 2 | 8.3200 | 0.0444 |
| HexNAc(5)Hex(4)Fuc(1) | G1FN/B | N[+1810] | 989.732 | 3 | >sp\|P01859\|IGHG2 | 8.3321 | EEQFN[8.987]STFR | 987 | 2 | 8.3200 | 0.0444 |
| HexNAc(4)Hex(4)Fuc(1)NeuAc(1) | G1FS | N[+1898] | 1019.060 | 3 | >sp\|P01859\|IGHG2 | 8.8296 | EEQFN[9.1012]STFR | 1012 | 2 | 8.9125 | 0.0155 |
| HexNAc(4)Hex(5)Fuc(1)NeuAc(1) | G2FS | N[+2060] | 1073.072 | 3 | >sp\|P01859\|IGHG2 | 8.8313 | EEQFN[9.1062]STFR | 1062 | 2 | 8.8259 | 0.0155 |

ND = not detected

**Supplementary Table S3. DIALib Y ion library for the eight cell wall glycopeptides with fixed RT at 12 min.**

**Supplementary Table S4. Peakview output of mesurement of ions and peptides using the Y ion library for the eight cell wall glycopeptides.**

**Supplementary Table S5. DIALib *by*, *by*+Y, stop*by*, and stop*by*+Y ion libraries for the eight cell wall glycopeptides with fixed RT at 12 min.**

**Supplementary Table S6. DIALib *by* ion library for the eight cell wall glycopeptides using published manually curated RTs [5].**

**Supplementary Table S7. DIALib stop*by* ion library for the eight cell wall glycopeptides using published manually curated RTs [5].**

**Supplementary Table S8. DIALib Y ion library for IgG glycopeptides across the full range of retention times and *m/z* windows.**

**Supplementary Table S9. Byonic search results for IgG *N*-glycopeptides.**

**Supplementary Table S10. ProteinPilot search results for cell wall glycopeptides from wild type yeast with potential *O*-glycosylation.**

**Supplementary Table S11. Byonic search results for cell wall glycopeptides from wild type yeast with potential *O*-glycosylation.**

**Supplementary Table S12. DIALib Y ion library for the Hsp150 cell wall T^318^SGTLEMNLK glycopeptide across the full range of retention times and *m/z* windows.**

**Supplementary Table S13. Peakview output of measurement of ions and peptides using the DIALib Y ion library from Supplementary Table S12.**

**Supplementary Table S14. DIALib oxonium ion library across the full range of retention times and *m/z* windows.**

**Supplementary Table S15. Peakview output of measurement of ions and peptides using the DIALib oxonium ion library from Supplementary Table S14.**

### Supplementary Figures

**Supplementary Figure S1.**
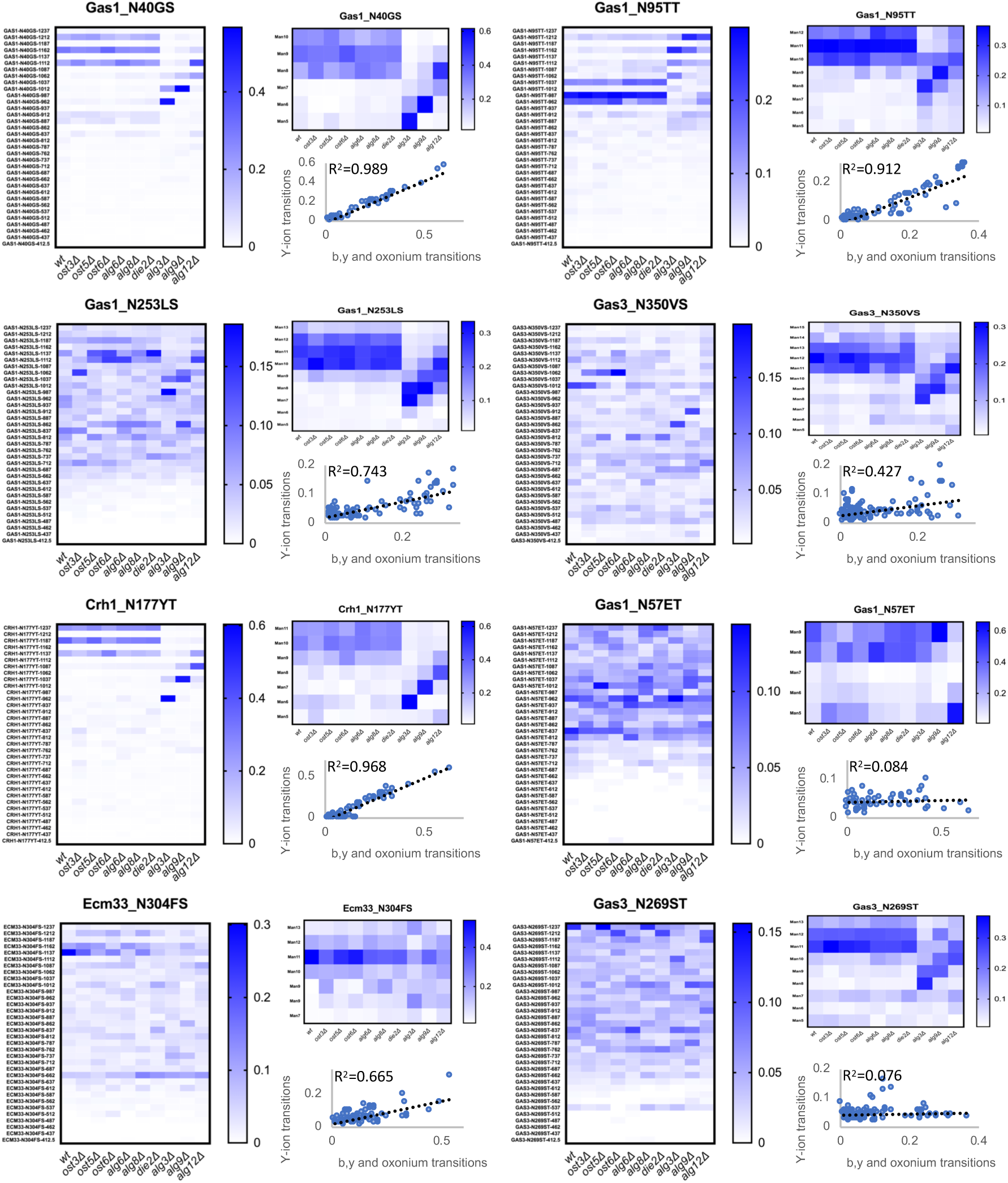
DIALib Y ion measurement of site-specific glycan structural heterogeneity for eight cell wall yeast glycopeptides. Abundance of glycoforms of glycopeptides containing Gas1 N^40^GS, Gas1 N^95^TT, Gas1 N^253^LS, Gas3 N^350^VS, Crh1 N^177^YT, Gas1 N^57^ET, Ecm33 N^304^FS, and Gas3 N^269^ST measured using a DIALib Y ion library or a manually curated ion library [5] in wild type yeast and yeast with mutations in the *N*-glycan biosynthesis pathway. Each square corresponds to the mean intensity of triplicate measurements. Also shown is the correlation between the data obtained using a DIALib Y ion library and a manually curated ion library [5] for each glycosylation site, with linear fit and R^2^ values.

**Supplementary Figure S2.**
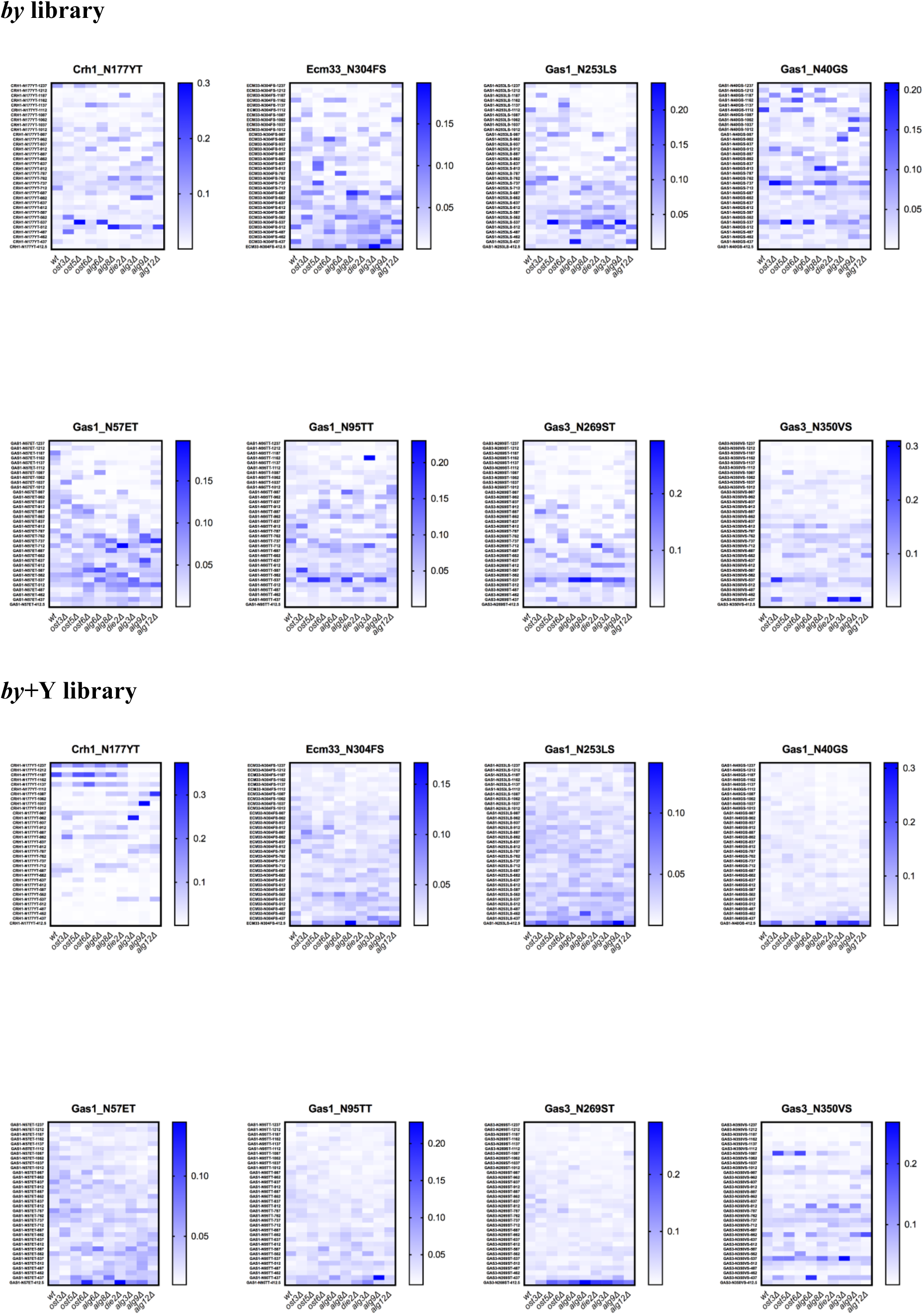
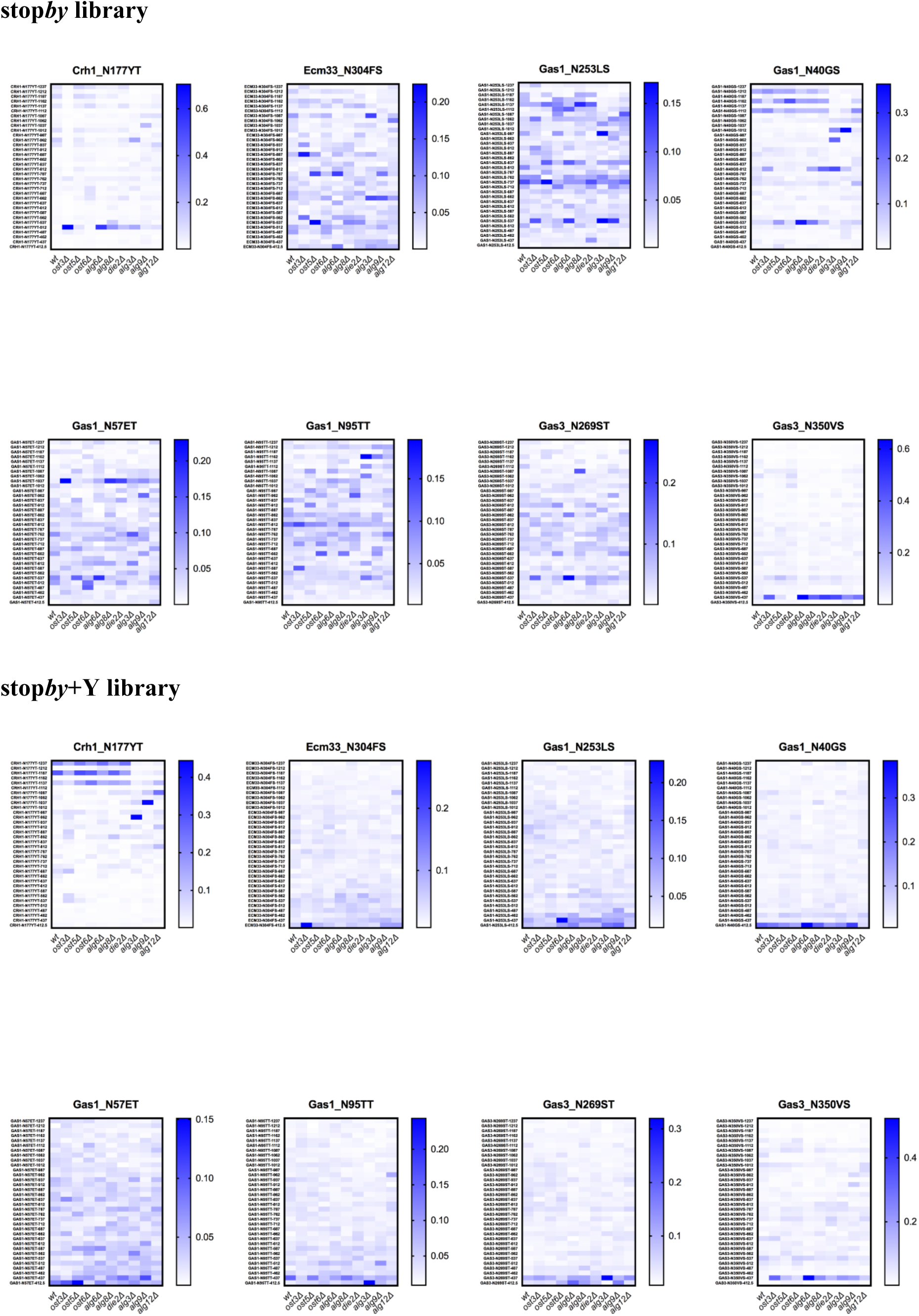
Theoretical DIALib *by*, *by*Y, stop*by*, and stop*by*Y ion libraries underperform compared to the DIALib Y ion library without optimization of peptide retention times. Abundance of glycoforms of eight cell wall glycopeptides measured using DIALib *by*, *by*Y, stop*by*, and stop*by*Y ion libraries in wild type yeast and yeast with mutations in the *N*-glycan biosynthesis pathway. Each square corresponds to the mean intensity of triplicates.

**Supplementary Figure S3.**
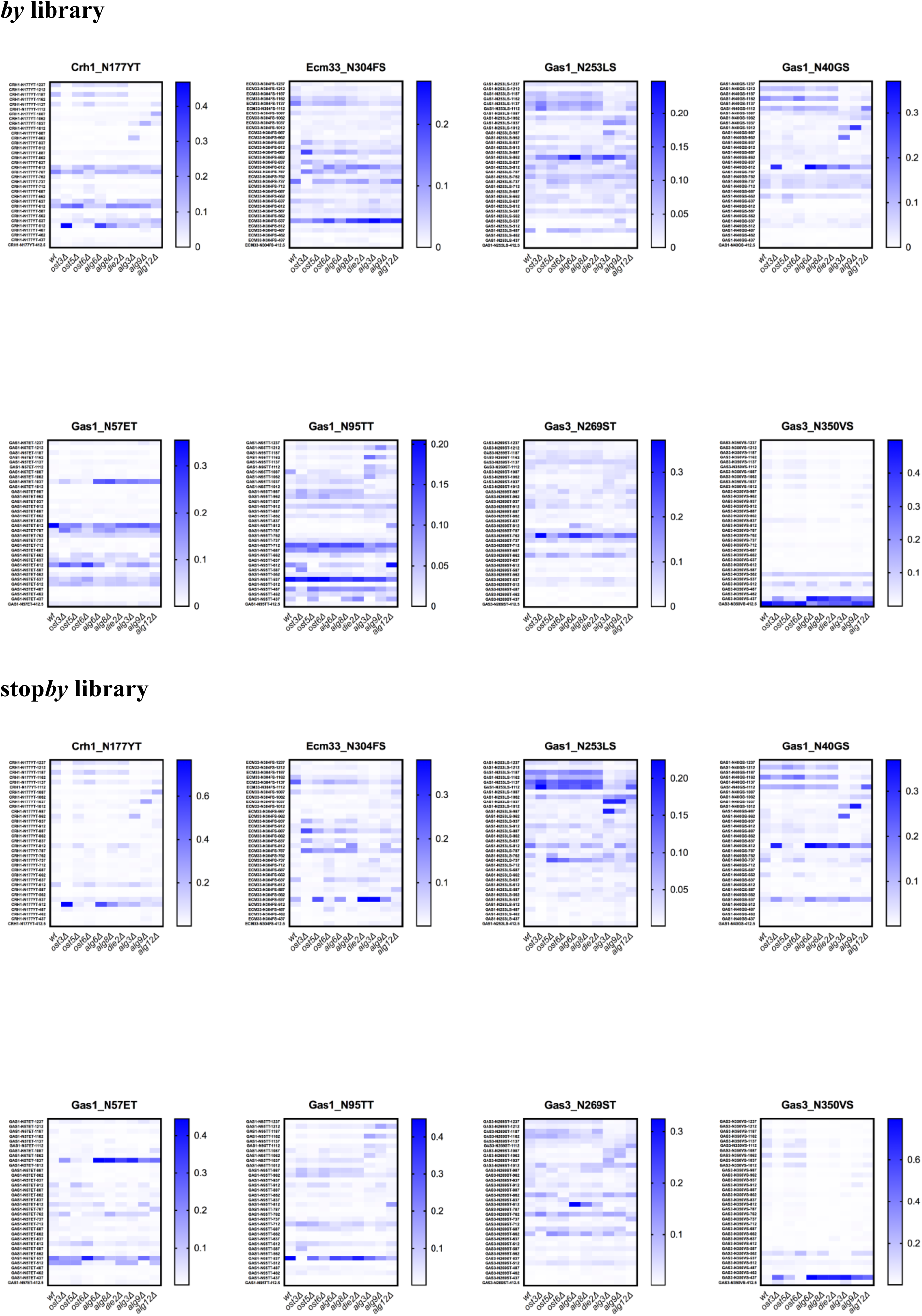
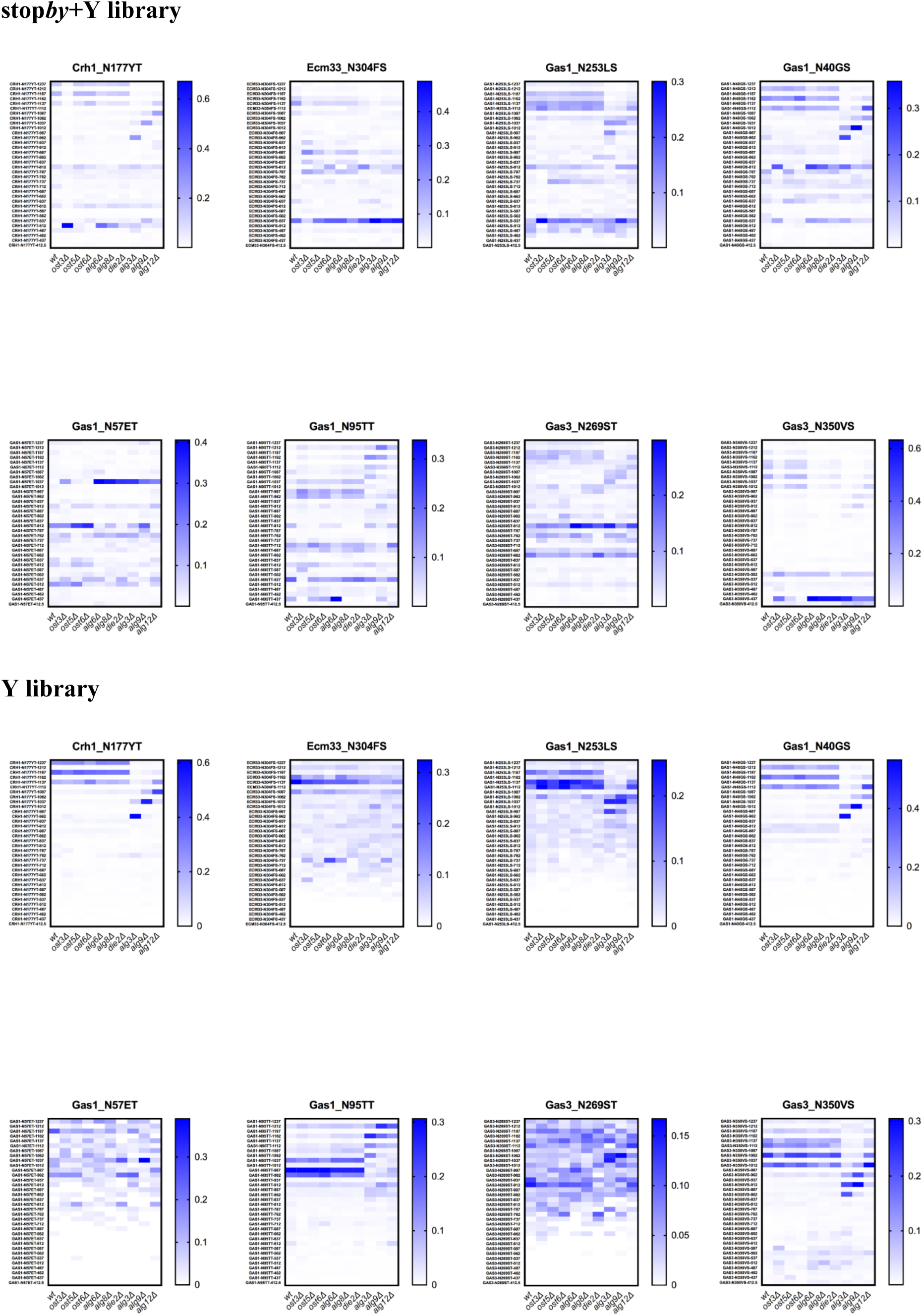
Optimized retention times improve peak picking for DIALib libraries. Abundance of glycoforms of all eight glycopeptides measured using the *by*, stop*by*, stop*by*+Y, and Y DIALib libraries and manually curated library incorporating experimental peptide retention times [5], measured in wild type yeast and in yeast with mutations in the *N*-glycan biosynthesis pathway.

